# Pairwise learning of MRI scans using a convolutional Siamese network for prediction of knee pain

**DOI:** 10.1101/463497

**Authors:** Gary H. Chang, David T. Felson, Shangran Qiu, Ali Guermazi, Terence D. Capellini, Vijaya B. Kolachalama

**Affiliations:** Section of Computational Biomedicine, Department of Medicine, Boston University School of Medicine, Boston, MA, USA – 02118; Section of Rheumatology, Department of Medicine, Boston University School of Medicine, Boston, MA, USA – 02118; Centre for Epidemiology, University of Manchester and the NIHR Manchester BRC, Manchester University, NHS Trust, Manchester, UK; Department of Radiology, Boston University School of Medicine, Boston, MA, USA – 02118; Department of Human Evolutionary Biology, Harvard University, Cambridge, MA, USA – 02138; Broad Institute of MIT and Harvard, Cambridge, MA, USA 02142; Whitaker Cardiovascular Institute, Boston University School of Medicine, Boston, MA, USA - 02118; Hariri Institute for Computing and Computational Science and Engineering, Boston University, Boston, MA, USA – 02215; Boston University Alzheimer’s Disease Center, Boston, MA, USA – 02118

**Keywords:** Osteoarthritis, Knee pain, Deep learning, MRI, Siamese network

## Abstract

**Objectives:** It remains difficult to characterize the source of pain in knee joints either using radiographs or magnetic resonance imaging (MRI). We sought to determine if advanced machine learning methods such as deep neural networks could distinguish knees with pain from those without it and identify the structural features that are associated with knee pain.

**Methods:** We constructed a convolutional Siamese network to associate MRI scans obtained on subjects from the Osteoarthritis Initiative (OAI) with frequent unilateral knee pain (n=1,529) comparing the knee with frequent pain to the contralateral knee without pain. The Siamese network architecture enabled pairwise learning of information from two-dimensional (2D) sagittal intermediate-weighted turbo spin echo slices obtained from similar locations on both knees. Class activation mapping (CAM) was utilized to create saliency maps, which highlighted the regions most associated with knee pain. The MRI scans and the CAMs of each subject were reviewed by an expert radiologist to identify the presence of abnormalities within the model-predicted regions of high association.

**Results:** Using 10-fold cross validation, our model achieved an area under curve (AUC) value of 0.808. When individuals whose knee WOMAC pain scores were not discordant were excluded, model performance increased to 0.853. The radiologist review revealed that about 86% of the cases that were predicted correctly had effusion-synovitis within the regions that were most associated with pain.

**Conclusions:** This study demonstrates a proof of principle that deep learning can be applied to assess knee pain from MRI scans.

## INTRODUCTION

Osteoarthritis (OA) is the most common musculoskeletal disease and one of the leading causes of disability globally [1]. The most incapacitating manifestation of OA is pain, and painful OA is most common in the knee. Severe OA-induced pain often leads to disability. Currently, there is no effective cure for advanced stage knee OA other than total joint replacement surgery. The occurrence of pain in knee joints with OA can be correlated with a variety of structural findings [2], and neuropathic mechanisms such as central sensitization and hyperalgesia [3–5]. For example, structural findings such as bone marrow lesions (BMLs), cartilage damage, synovitis, and effusion are related to pain in knee joints with OA [2, 6–9]. Also, the frequency and severity of pain are self-reported and usually defined subjectively [3, 10]. Consequently, the correlation between pain and radiographic findings is weak and there has been little success in correlating OA-induced pain with a specific type and location of structural damage. One review found that the proportion of subjects with knee pain who have radiographic OA ranged from 15 to 76% [11]. Identifying the source and location of OA-induced pain in each individual could greatly benefit the design of targeted, individualized treatments to reduce symptoms and to limit disability [10]. Further, for those with knee pain as part of a widespread pain diathesis, determining the absence of pain inducing knee pathology might aid in diagnosis.

MRI scans are capable of providing more detailed structural information about the knee joint than radiographs. A systematic review of MRI measures found that knee pain may arise from BMLs, effusion, and synovitis; however, the correlation between pain and MRI findings was inconsistent and moderate at best. [12–14]. Further, since MRI findings associated with pain are often present in multiple locations in the knee and since many knees without pain also have MRI findings, the lesions associated with knee pain are hard to localize and identify. This limits treatment approaches that seek to treat localized areas of the knee including surgeries such as unicondylar replacements and cryoneuroablation procedures, and hampers the development of targeted treatment strategies. Further rehabilitation strategies which focus on lessening the load to painful regions of the knee are limited. Hence, there is a need to develop a method to objectively and accurately associate MRI scans with knee pain. The first step of such an approach is to identify MRI findings that discriminate painful from nonpainful knees and then to determine the regions within the knee that are the likely source of pain. In this paper, we pursue the first step, using machine learning (ML) approaches to discriminate between painful and nonpainful knees. We begin to address the second step by identifying lesions identified in the first step as potential sources of pain.

ML is a discipline within computer science that uses computational algorithms for the analysis of various forms of data. These algorithms applied to medical images have shown remarkable success in predicting various outcomes of interest [15, 16]. Over the past few years, a new ML modality known as deep learning is gaining popularity because of its ability to analyze large volumes of data for pattern recognition and prediction with unprecedented levels of success. Specifically, deep learning frameworks such as convolutional neural networks (CNN) are increasingly being leveraged for object recognition tasks and specifically for disease classification [17]. Traditional ML algorithms require visual or statistical features to be manually identified and selected (“handcrafted”), and researchers need to decide which of these handcrafted features are related to the problem at hand. On the contrary, deep learning algorithms extract visual features automatically, and one can utilize them simultaneously for various applications related to classification, segmentation, and detection [15]. CNN model training is associated with learning a series of image filters through numerous layers of feed-forward neural networks. The filters are then projected on the original input image, and the image features that are most correlated with the outcome are extracted through the training process. Recently, deep learning techniques were applied on MRI scans of the knee joint for automatic cartilage and meniscus segmentation [18], and detection of ligament and meniscal tears [19], and on x-ray imaging for automatic Kellgren-Lawrence (KL) grade classification [20].

The purpose of this study was first, to investigate the performance of a deep learning framework to differentiate painful knees from nonpainful ones and second to identify the structural lesions that are most relevant to knee pain using MRI scans of both knees of individuals enrolled in the Osteoarthritis Initiative (OAI). Unilateral frequent knee pain was defined when an individual had pain, aching or stiffness for more than half of the days in a month in one knee and no pain in the contralateral knee. We used sagittal intermediate-weighted turbo spin echo (SAG-IW-TSE) sequence images that capture structural regions thought to be critical in generating knee pain, BMLs, synovitis, effusion and cartilage loss [6, 21–26], to train a convolutional Siamese network. We subsequently leveraged class activation mapping (CAM) to identify the structural regions that were most relevant to frequent knee pain. An expert radiologist then independently reviewed the MRI scans and identified possible presence of knee abnormalities, which were then compared with CAM-based findings.

## METHODS

### Study selection

Our samples were drawn from the OAI, an NIH funded study of persons with or at risk of knee OA [27, 28]. Baseline data from the OAI database was used for training and testing our deep learning model (**Table 1**). The dataset consists of MRI scans and clinical data from 4,796 subjects, of whom 1,606 subjects had pain, aching, or stiffness for more than half the days of a recent month in one knee and who reported they did not have pain on more than half the days in the contralateral knee. Out of them, 781 subjects had frequent knee pain in the left knee (label 0), and 825 subjects had frequent knee pain in the right knee (label 1). Out of these cases, we found 1,529 subjects that had the MRI sequence (SAG-IW-TSE) available in both knees. From these cases, 1,505 subjects passed initial quality check (see below), and were used for construction of the deep learning model (Model A). Additionally, we excluded subjects with similar Western Ontario and McMaster Universities Osteoarthritis Index (WOMAC) pain scores between the left and right knee to construct another model. For this case, we selected a subset of subjects who had a WOMAC pain score difference greater than 2 between the knees. In total, 721 subjects met the criteria and 710 subjects were used for model construction after quality check (Model B). For the selected subjects, sagittal intermediate-weighted turbo spin echo (SAG-IW-TSE) sequence images were extracted as the inputs and unilateral knee pain as the output to solve a binary classification problem using the deep learning framework. The SAG-IW-TSE sequence of OAI images provided great details in the knee including bone marrow abnormality, synovial effusion, osteophytes, and cysts [27]. The original 2D MRI slices had dimensions of 480×448 pixels. The imaging was performed with a 3.0 T magnet using imaging sequence with TR/TE of 3200/30 ms. The in-plane spatial resolution was 0.357×0.511 mm and the slice thickness was 3.0 mm [27].

### Image registration and quality check

The majority of the SAG-IW-TSE sequence scans comprised 37 slices per knee. We first manually examined all the MRI slices oriented in the sagittal direction and selected the slice showing the most complete view of posterior cruciate ligament (PCL) and indexed it as the center slice (Red colored box in **Figure 1A**). The remaining slices were indexed relative to the center slice for each knee, with a plus sign indicating the lateral direction and a minus sign indicating the medial direction. After the images were indexed, for each 2D MRI slice of the knee for a subject, we performed Euclidean transformation to align the slice with respect to a template that we previously picked from the MRI scans of a subject (**Figure 1B**). A region with dimensions of 294×294 pixels (105×105 mm) was subsequently selected, which contained the femoral, tibial, and patellar components of bone, cartilage and meniscus and all the registered slices were cropped to this region of interest. All the cropped slices were further resized into images of 224×224 pixels, and 11 adjacent slices on the lateral side of the center slice, 11 adjacent slices on the medial side of the center slice, and the center slice were selected for deep learning. In total, 46 MRI slices were selected for each subject (23 from each knee). After image registration, we performed a quality check that involved manually observing each sagittal slice of each scan for artifacts such as missing data, abnormal misalignment within a slice, presence of a foreign object, and cases with sub-regions of high contrast. All these cases were not considered for model training or testing (**Figure 1C**).

**Figure 1:**
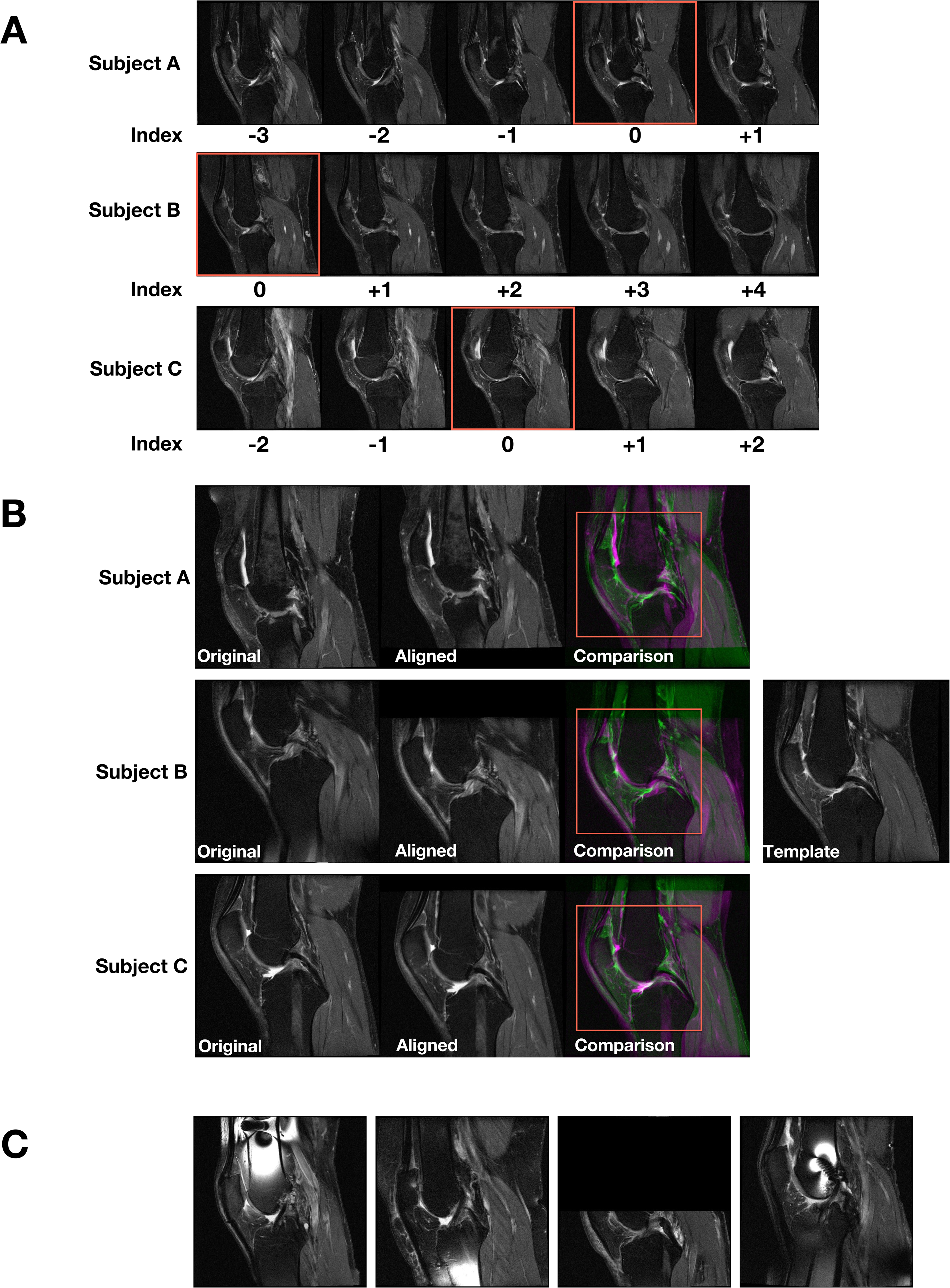
Image processing pipeline. **(A)** For each subject’s knee joint, we manually examined all the MRI slices oriented in the sagittal view and selected the slice showing the posterior cruciate ligament (PCL) and indexed it as the center slice (red colored box). The remaining slices were indexed relative to the center slice for each knee. Two-dimensional MRI slices for three different subjects are shown. **(B)** For each 2D MRI slice of the knee for a subject, we performed linear registration to align the slice with respect to a template that was already selected after manual examination. Later, a region (red colored box) containing the center of the knee joint with dimensions 294×294 pixels was cropped for all registered slices and used for model training. Three cases from the baseline OAI dataset are shown. **(C)** Sample cases not used for model training due to the presence of various artifacts.

### Neural network architecture

We developed a deep neural network inspired by the Siamese network architecture [29], to associate the MRI scans from the left and right knee of the same subject with knee pain. Siamese networks were originally proposed to estimate similarities between a pair of images, and it was accomplished by feeding each of the two input images to two identical neural networks that shared the same parameters. The outputs from the two branches were subsequently joined by a contrastive loss function. In the present study, our model architecture was designed such that a pair of MRI slices from the two knees, each extracted from a relatively similar location within each knee were learned together **(Figure 2A)**. The deep learning architecture consisted of convolutional, batch normalization, nonlinear activation and two max pooling layers.

**Figure 2:**
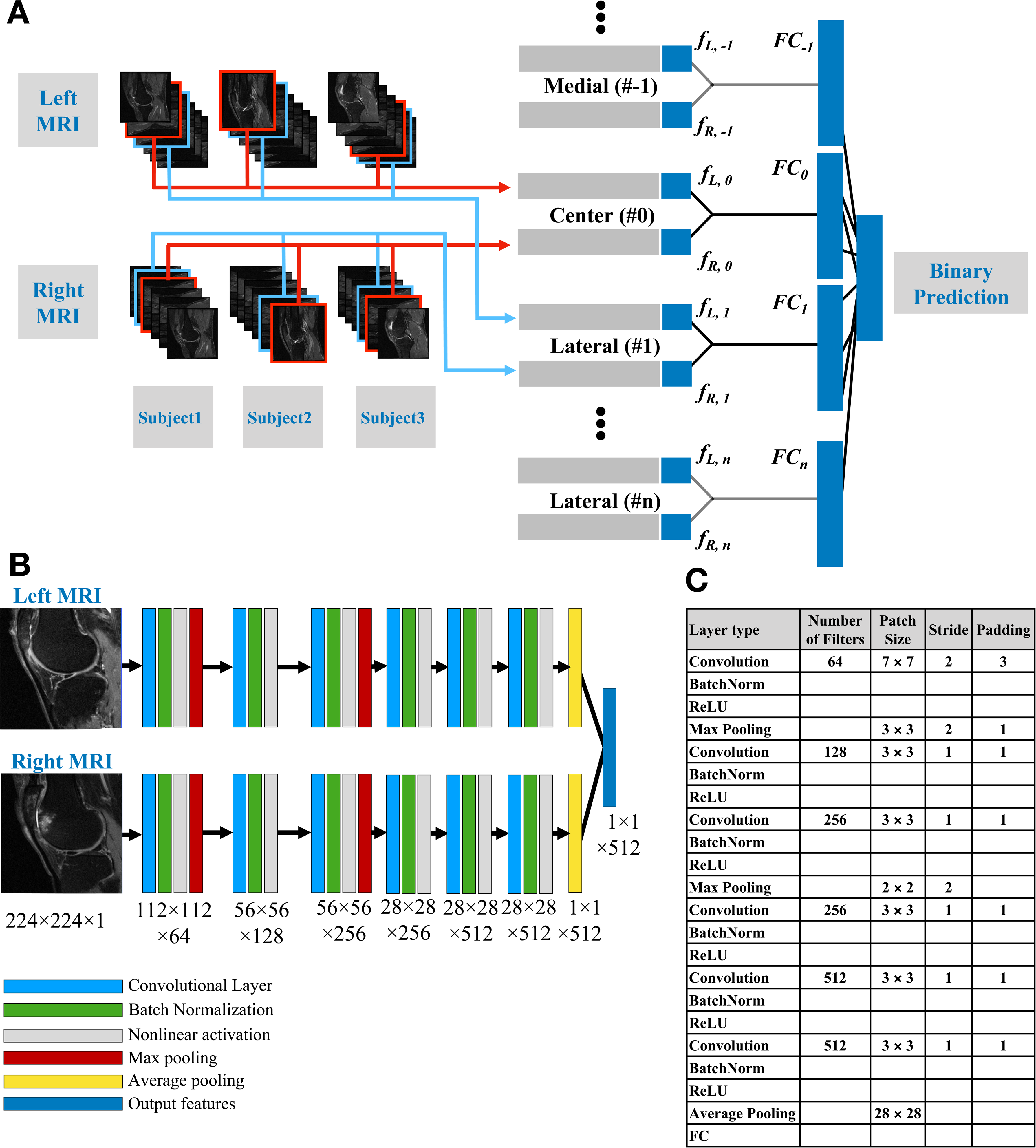
Architecture of the convolutional Siamese network. **(A)** After imaging pre-processing, the indexed MRI slices were fed into the corresponding Siamese network model. The outputs from all the 2D models were concatenated for the binary prediction task. **(B)** The 2D MR slices for the left and right knee were fed into the network simultaneously. A series of convolutional operations, batch normalization, nonlinear activation, max pooling and average pooling were applied to predict knee pain, by solving a binary classification problem.

Each neural network that is part of the Siamese architecture comprised of 6 convolutional layers (**Figure 2B**). For layers 1 and 3, each convolutional operation was followed by batch normalization, nonlinear activation and max pooling, whereas for layers 2, 4, 5, and 6, each convolutional operation was followed by just batch normalization and nonlinear activation. Only the first convolutional layer and the two max pooling layers had a stride of 2, whereas the other layers had a stride of 1. Consequently, the final convolutional layer had a high in-plane resolution with a dimension of 512×28×28 **(Figure 2C)**. It is worthwhile to note that while our network was a *de novo* design, it has some similarities to the original VGGNet architecture [30].

During the training process, the two branches of the Siamese network learned the same set of shared parameters by presumably focusing on the similarities as well as the unique aspects of each knee associated with pain. This also allowed us to reduce the number of learnable parameters, considering both the left and right knee were evaluated by the same architecture. The training was performed using stochastic gradient descent optimizer and cross-entropy loss function for binary classification. A batch size of 64 was used and the images were normalized between 0 and 1 before training. The 2D models was trained with an initial learning rate of 1e-4 and the learning rate decayed by a factor of 0.1 every 20 epochs of training. The model converged after 500 epochs of training using stochastic gradient descent algorithm with momentum of 0.9. The output features from all the 2D models were subsequently concatenated for the binary prediction task. The neural network model, optimizer and training algorithms were implemented using an open-source deep learning library (PyTorch). Model training was performed on a high-performance workstation containing 4 NVIDIA GPUs (RTX 2080Ti), and the training time was about 13.6 hours for Model A and about 8.7 hours for Model B.

### Class activation mapping and radiologist review

We used the technique of class activation mapping (CAM) to generate saliency maps that highlighted regions that were most associated with pain, at least as determined by the neural network. CAMs have the ability to localize the discriminative image regions from CNN models trained for classification without any prior locational knowledge [31]. This was achieved by first extracting the final feature maps of the left and right knee of the subject (*f_L,n_* and *f_R,n_* in **Figure 2A**, respectively). These feature maps were generated by the final convolutional layer of the network and had a dimension of 512×28×28. The maps were subsequently multiplied by weights of their respective fully-connected layer (*FC_n_* in **Figure 2A**), which indicated the importance of each feature stored in the extracted maps. This resulted in CAMs with a dimension of 28×28 pixels for each MRI slice. For each subject, we identified a CAM with the most pain-relevant regions by selecting the CAM with highest average value from the 23 CAMs generated from all the MRI slices. This process resulted in the generation of saliency maps that highlighted the structural regions that were most associated with unilateral knee pain.

The dataset of 710 subjects (unilateral knee pain, between knee WOMAC pain difference **≥** 3) was divided into training, validation and testing sets in a 70:15:15 ratio (**Table 2**). The sampling was stratified based on risk factors of OA, including gender, age and body mass index (BMI). A musculoskeletal radiologist with extensive experience in knee MRI interpretation reviewed CAMs of MRI of the last 15%, consisting of 107 subjects. The radiologist reviewed each case and identified the presence of abnormalities using the MRI scan. The radiologist then reviewed the model-derived CAMs and identified the specific lesion that was co-localized with the highlighted region in CAM.

### Model performance and statistical analysis

To evaluate model performance, we first performed 10-fold cross-validation on the model using the selected dataset of 1,505 subjects (Model A, unilateral knee pain) and 710 subjects (Model B, unilateral knee pain, WOMAC pain difference **≥** 3) (**Table 2**). In each fold, the selected dataset was divided into the training and testing sets in a 9:1 ratio. In this scenario, every subject in the entire selected dataset appeared in the testing set exactly once. To evaluate the effectiveness of image pre-processing, we trained two separate models without image registration by simply selected 23 slices (8-30th) in the center of the MRI scans before alignment as input images. We subsequently computed the area under curve (AUC) of the receiver operating characteristic (ROC) curves of the binary classifier.

Descriptive statistics are presented as the mean along with the 95% confidence intervals. Unpaired Student’s t-test was used to compare the mean value of two different groups, and Fisher’s exact test was used to examine the non-random association between two groups of categorical variables. A p-value < 0.01 was considered statistically significant.

## RESULTS

Among the 1,505 subjects from the baseline OAI (n=4,759) (Model A in Tables 1a & 1b), the mean age was 60.7±9.1 years and the mean BMI was 28.7±4.7 kg/m^2^. About 56.9% of the selected subjects were women. When cases with WOMAC score difference <3 were excluded (Model B in Tables 1a & 1c), the sub-group demographic characteristics remained relatively similar with respect to the overall group (Mean age: 60.9±9.2 years; Mean BMI: 29.2±4.8 kg/m^2^ and Percentage women: 60.1). For the cases that were reviewed by the radiologist (Table 1d), stratified sampling based on age, BMI, and gender helped us to generate a sub-group with demographic characteristics that were similar to the ones considered for models A & B.

**Table 1:** Schematic of study selection. (Table 1a): In total, 1,606 subjects with unilateral frequent knee pain were selected from the OAI’s baseline study, consisting of 4,796 subjects. After image quality check, 1,505 subjects were selected to train the neural network (Model A). Subjects who had the contralateral difference in WOMAC pain score (≥ 3) were further selected, leading to 710 subjects which were then used to train another neural network (Model B). Stratified sampling was then used to divide the dataset in the ratio of 70:15:15, where 70% of the dataset was used for training, 15% for validation, and the remaining cases (n=107) for independent testing. The stratified sampling was performed to match for age, gender, and body mass index. These cases were then reviewed by the expert radiologist. (Tables 1b-1d): Baseline characteristics of the sub-group of individuals selected for model training (Models A & B), and for radiologist review.

|  |  |  |  |  |
| --- | --- | --- | --- | --- |
| Number of subjects in OAI baseline dataset | Model A (n = 1505) |  |  |  |
| n = 4796 | Parameter | Left Knee Pain | Right Knee Pain | p-value |
| Cases with unilateral knee pain | Subjects | n = 734 | n = 771 |  |
| n = 1606 | Percent female | 57.9 | 55.9 | 0.700 |
| Unilateral knee pain cases selected after quality check (Model A) | Age (years) | 60.13 ± 9.18 | 61.24 ± 8.98 | 0.018 |
|  | BMI (kg/m2) | 28.97 ± 4.88 | 28.38 ± 4.53 | 0.015 |
| n = 1505 |  |  |  |  |
| Unilateral knee pain cases with contralateral WOMAC pain score difference (≥3) (Model B) | Model B (n = 710) |  |  |  |
|  | Parameter | Left Knee Pain | Right Knee Pain | p-value |
| n = 710 | Subjects | n = 350 | n = 360 |  |
| Cases selected for radiologist review | Percent female | 59.4 | 60.8 | 0.854 |
| n = 107 | Age (years) | 60.62 ± 9.39 | 61.21 ± 9.01 | 0.395 |
|  | BMI (kg/m2) | 29.50 ± 4.96 | 28.94 ± 4.63 | 0.120 |
| MRI scan type | Cases for radiologist review (n = 107) |  |  |  |
| SAG_IW_TSE_RIGHT, SAG_IW_TSE_LEFT | Parameter | Left Knee Pain | Right Knee Pain | p-value |
| 2D MRI slice size | Subjects | n = 46 | n = 61 |  |
| 448×444 pixels | Percent female | 60.9 | 59.0 | 1.000 |
|  | Age (years) | 60.83 ± 9.34 | 60.67 ± 8.90 | 0.932 |
|  | BMI (kg/m2) | 29.38 ± 5.30 | 29.01 ± 4.89 | 0.711 |

Our methodology of generating CAMs provided a means by which to identify and examine the regions that were highly associated with knee pain. These CAMs are generated by extracting the features learned from the final convolutional layer in the neural network. Therefore, the ability of the CAMs to precisely identify a region of interest is directly dependent on the spatial resolution of this convolutional layer. Previously, researchers who proposed CAMs have used model architectures that had an in-plane resolution of 14×14 pixels (**Figure 3A**) [31]. In the present study, we generated CAMs with higher resolution containing 28×28 pixels, which resulted in qualitatively improved identification of the regions associated with pain (**Figure 3B**). For comparison and to evaluate statistical significance, we performed binary thresholding of the CAM segmented areas for both cases. When a threshold of 0.5 was used on the CAMs, we found that only 6.0±2.4% of the overall image area was segmented from the CAMs generated by our model. In comparison, 22.3±9.4% of the image area was segmented when the same threshold was used on the CAMs generated by the model used by Zhou and colleagues [31].

**Figure 3:**
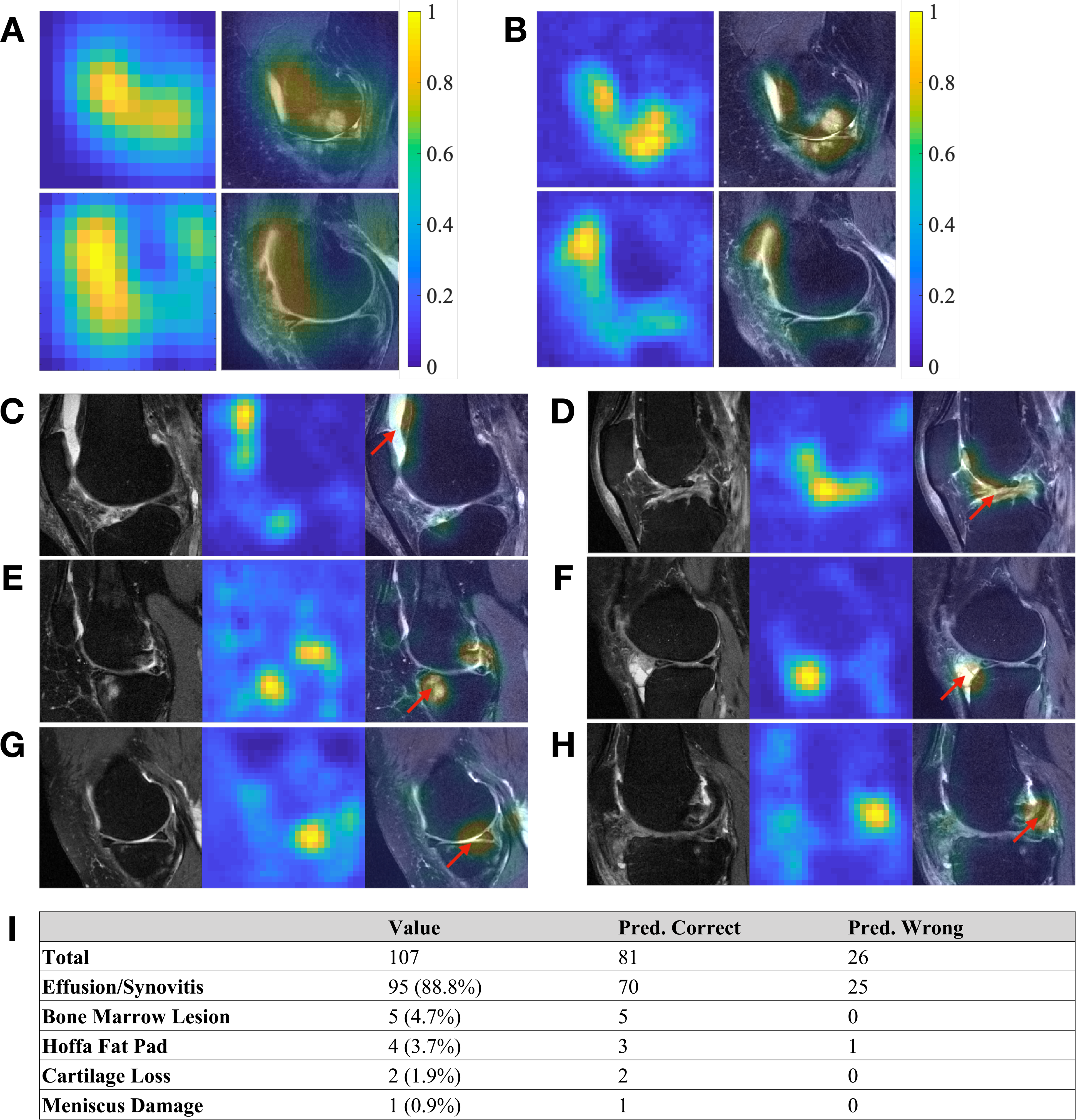
Generated CAMs on selected subjects within the test data. **(A)** Examples of CAMs generated from the fine-tuned VGGNet model, resulting in CAMs with an in-plane resolution of 14×14 pixels. **(B)** Examples of CAMs generated from the present model from the same MRI images with an in-plane resolution of 28×28 pixels. Both the heat maps and the overlap of the MR image with the heat map are shown. In some cases of the test data **(C, D)**, effusion/synovitis was identified as the lesion present within the hot spots. Also, in few other cases, **(E)** bone marrow lesions, **(F)** Hoffa fat pad abnormality, **(G)** cartilage loss, and **(H)** meniscus damage, were identified as the lesions present within the hot spots. The red arrows indicate the locations of the identified structural regions. **(I)** Radiologist’s assessment on the test cases (n=107). For each case, the MR scan and the model-derived heat map of the knee with confirmed pain were reviewed by the radiologist, who then identified the presence of any lesions within the regions highlighted by the heat map.

After the radiologist review, the location and the type of lesions that were co-localized with the highlighted region in the CAMs of subjects were identified (**Figures 3C-H**). The identified lesions included effusion, synovitis, BML, Hoffa fat pad lesion, cartilage loss, and meniscal damage. The effusion on the selected intermediate-weighted MRI scans included effusion and synovitis, and therefore, we combined effusion/synovitis into a single category, as used in MRI Osteoarthritis Knee Score (MOAKS). Out of the 107 cases reviewed by the radiologist (**Figure 3I**), effusion/synovitis was found to be the most relevant structural abnormality related to frequent knee pain in 95 (88.8%) subjects. BML was found to be the most relevant abnormality for 5 (5.6%) subjects. Hoffa fat pad abnormalities were found for 4 (3.7%) subjects, cartilage loss was found for 2 (1.9%) subjects, and meniscal damage was found for 1 subject.

The overall performance of the model was evaluated using 10-fold cross validation. For the model alone not including age, sex or other variables trained with 1,505 subjects with unilateral knee pain (Model A), we observed an AUC of 0.808 on the test data (**Figure 4A**). For the case with 710 subjects with unilateral knee pain and with a difference in WOMAC pain score larger than 2 (Model B), the model achieved an AUC of 0.853 on the test data (**Figure 4B**). In comparison, models trained without image registration achieved AUCs of 0.769 and 0.812 on Model A and Model B, respectively.

**Figure 4:**
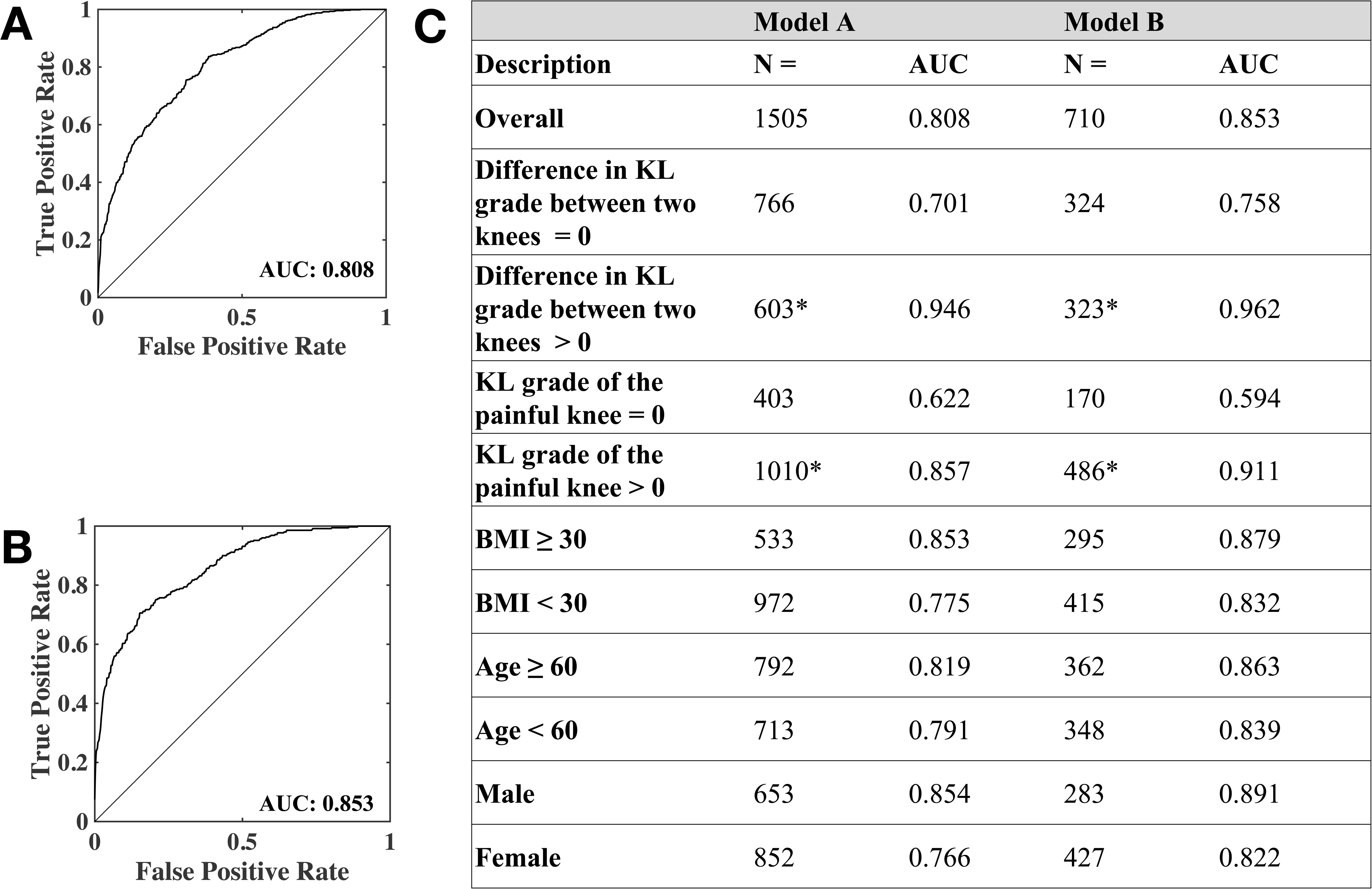
Performance of the convolutional Siamese network model. (A) Receiver operating characteristic (ROC) curve of Model A trained with 1,505 subjects with unilateral frequent knee pain. (B) ROC curve of Model B trained with 710 subjects with unilateral frequent knee pain and contralateral knee difference in WOMAC pain ≥ 3. (C) Sub-group analysis. Performance of the models as a function of KL grade, BMI, age, and gender. The asterisk (*) is used to indicate that KL-grade was not available on few cases for the sub-group analysis.

Sub-group analysis further revealed that model performance varied as a function of KL-grade, BMI, age, and gender (**Figure 4C**). An interesting finding from our study is that our model could differentiate painful from nonpainful knees even when the OA disease stages were similar between the two knees (i.e. Difference in KL grade = 0, AUC=0.701 for Model A and AUC=0.758 for Model B, **Figure 4C**). However, when there was a difference in radiographic OA between the knees (i.e. Difference in KL grade ≠ 0), then the model performance increased by about 26% for Model A and about 21% for Model B (**Figure 4C**). Our study also revealed that when the KL-grade of the painful knee was 0, then the model resulted in a modest performance (AUC=0.622 for Model A and AUC=0.594 for Model B, **Figure 4C**). When the KL-grade was > 0, the model was able to better distinguish between the painful knee from the non-painful knee (Performance increased by 27% for Model A and about 35% for Model B, **Figure 4C**). It is also worthwhile to note that the WOMAC pain score of the painful knee was significantly lower for KL=0 cases than the KL>0 cases (3.10 vs 4.20, p ≪ 0.01). The models also had higher values of AUC for subjects who were older, male and had higher BMI (**Figure 4C**). Specifically, we observed that subjects age 60 and older had a higher average KL-grade in the painful knee than the subjects who were younger than 60 years (1.71 vs 1.24; p<0.01). Also, males had higher averaged KL-grade in the painful knee than females (1.62 vs 1.37; p<0.01). Subjects with BMI greater than 30 also had higher averaged KL-grade in the painful knee than the subjects with lower BMI than 30 (1.8 vs 1.31; p ≪ 0.01).

## DISCUSSION

We developed a deep learning approach to distinguish painful knees from nonpainful contralateral ones, and we were able to do so with a high level of accuracy. Further, an expert radiologist reviewing the areas within the knees that were identified as painful suggested that these were primarily regions of synovitis or effusion, which are known sources of knee pain in OA. Several studies have examined the association between radiographic features in the knee and knee pain within individual subjects and across multiple cohorts [13, 32–34]. While some studies relied on x-ray imaging, other studies relied on more sophisticated modalities such as MRI scans [35]. For some cases, associations between imaging features and unilateral pain were observed by comparing the knee with pain in the individual with the contralateral knee without pain in the same individual [36].

The Siamese network that used both knees for the same individual to study unilateral pain effectively allowed one knee to serve as a control to the other knee with pain. Moreover, this dual-knee paradigm to assess knee pain is an attractive choice because the effect of common confounding factors such as age, gender, and BMI, on the outcome of interest no longer applies in such scenarios. Also, unlike the manual extraction of image-based or radiographic features which were then associated with knee pain, we investigated the feasibility of using deep learning to correlate structural regions from MRI scans of both knees with unilateral frequent knee pain.

By combining information from a series of 2D slices, our model synthesized needed information from multiple locations to predict knee pain. This strategy resulted in a model that achieved high AUC (0.808), as evaluated using 10-fold cross validation, and model performance improved by about 5.6% when subjects with similar WOMAC pain score between the two knees were excluded (AUC=0.853). This improvement suggests that the Siamese network was able to identify the image features that were associated with a strong pain signature arising from one knee. This also implies that a set of image features that are common between the knees was also learned, but were not considered by the model to play a role in predicting knee pain. When compared to a machine learning approach using postero-anterior and lateral knee x-rays to predict knee pain, our model generated a significantly higher AUC in predicting unilateral knee pain [37]. We do however acknowledge that the training and testing datasets used for this reported study were different than the dataset presently used, and therefore a head-to-head comparison between the results is not feasible.

Our model’s improved performance when the difference in KL-grade was non-zero (**Figure 4C**), underscores the notion that prevalence and severity of knee pain is greatly influenced by the presence of pre-radiographic OA (KL=1) or radiographic OA (KL>1). This may imply that the pain symptoms and potential structural abnormalities are perhaps less severe than the knees with frequent knee pain with pre-radiographic or radiographic OA. Indeed, previous studies have reported that knees of KL grade 4 were 73-151 times more likely to have pain than knees of KL grade 0 [35].

While deep learning algorithms such as CNNs are now increasingly considered for image-based classification, most of the previous work focused on using datasets containing single 2D MRI slices or radiography images converted to 2D grayscale formats. An important reason for using such data is because developing a three-dimensional (3D) deep neural network would require higher amounts of GPU memory as more parameters need to be learned to fully train the network. To circumvent this problem, we extracted features from each 2D MRI slice and then combined them using fully-connected layers and then associated them with the output of interest. This framework therefore served as the best compromise between using as much information as possible from the volumetric data available within an MRI scan and the ability of utilizing such datasets to train without causing memory-related errors.

The series of pre-processing steps, including manual quality check and image registration of each MRI slice, helped our model focus on a region representing the knee joint. With the network architecture designed to preserve high in-plane resolution in the CAM, we were able to extract slice-specific CAMs from each subject. Using a straightforward Euclidean transform, we aligned all the slices with respect to a pre-defined template, and then isolated a region representing the knee joint from all the registered images. More sophisticated linear or even nonlinear image registration techniques may also be applied to improve the alignment of structures of interest between different subjects. Nonetheless, it is important to note that certain forms of nonlinear registration techniques may introduce unwanted distortion, which then may lead to invalid representations of important anatomical features [38].

For the present study, we trained two identical neural networks in the ‘Siamese’ sense to generate a model that led to prediction of knee pain with high accuracy. In principle, pre-trained neural networks (i.e. VGGNet, AlexNet, etc.) could be used as part of the Siamese architecture. In order to evaluate this aspect, we generated another Siamese architecture using the network proposed by Zhou et. al. [31] (VGG-GAP), as the neural network for pairwise learning. We used the data split that was used for the radiologist review to train this model (**Table 2**). Interestingly, the AUC value on the test data for this case was 0.856. We also trained another Siamese architecture using a modified version of the AlexNet (AlexNet-GAP [31]) as the neural network for pairwise learning. For this case, the AUC value on the test data was 0.863. In comparison, our proposed *de novo* Siamese network resulted in an AUC value of 0.862 on the test data.

**Table 2:** Data partitioning and modeling. Model A and Model B were constructed on the 1,505 subjects and 701 subjects, respectively. Performance of these models was evaluated after 10-fold cross validation. For radiologist review, we split the 710 cases in 70:15:15 ratio using stratified sampling. A new model was trained using 70% of the data and 15% of the data was used for internal validation. The remaining 15% was used for testing and radiologist review.

| Model A<br>(Unilateral knee pain) | 10-Fold Cross Validation |  |  |
| --- | --- | --- | --- |
|  | Training | Validation | Testing |
| n = 1505 | 90% | NA | 10% |
| Model B<br>(Unilateral knee pain, WOMAC pain difference ≥ 3) | 10-Fold Cross Validation |  |  |
|  | Training | Validation | Testing |
| n = 710 | 90% | NA | 10% |
| For radiologist review | Stratified Sampling |  |  |
|  | Training | Validation | Testing (for review) |
| n = 710 | 70% (496) | 15% (107) | 15% (107) |
| Percent female | 59.2 | 62.0 | 60.1 |
| Age (years) | 61.0±9.3 | 60.5±9.0 | 6.06±9.1 |
| BMI (kg/m2) | 29.1±4.7 | 29.4±5.0 | 29.2±5.0 |

Results from these models imply that our proposed Siamese network achieved performance values that were similar to Siamese architectures based on pre-trained networks. However, the major difference is that our proposed network had high spatial resolution in the CAMs to allow us to associate the results of CAMs with specific anatomical regions, and consequently to identify lesions that were highly correlated with knee pain.

While our deep learning model demonstrates promising results for predicting unilateral knee pain using MRI scans, there is room for improvement in model performance. Other neural network architectures can be explored such as deep autoencoders [39]. In a recent examination of x-rays and their prediction of knee pain, limiting the pain outcome to subjects who repeatedly reported knee pain increased the accuracy of x-ray prediction. Alternate definitions of pain or tenderness could facilitate development of models with higher performance. Our fusion model framework used a series of sagittal MRI slices, and it has the capability to combine coronal and axial MRI slices as well.

In conclusion, this work demonstrates the use of a convolutional Siamese network that simultaneously associated MR imaging data of both knees with unilateral knee pain. This framework allowed us to combine data from multiple 2D MRI slices from both the knees to efficiently construct the deep learning model. Such a modeling strategy can be easily extended to predict other clinical outcomes of interest. Our results provide a means by which to understand and evaluate early imaging markers of OA and other joint disorders. Further validation of the deep learning model across different imaging datasets is necessary to validate this technique across the full spectrum of OA.

## CONTRIBUTORSHIP

GHC designed the neural network and performed the analysis; GHC, DTF, TDC and VBK conceptualized the study; SQ assisted data analysis; AG reviewed the MRI scans; GHC and VBK wrote the manuscript; DTF, AG and TDC edited the manuscript; VBK directed the entire study.

## DATA SHARING INFORMATION

Data and computer scripts are available on GitHub (https://github.com/vkola-lab/mrm2019).

## ETHICAL APPROVAL INFORMATION

Not applicable

## PATIENT AND PUBLIC INVOLVEMENT

This research was done without patient involvement. Patients were not invited to comment on the study design and were not consulted to develop patient relevant outcomes or interpret the results.

